# Resting-state functional connectivity of social brain regions predicts motivated dishonesty

**DOI:** 10.1101/2021.11.10.468161

**Authors:** Luoyao Pang, Huidi Li, Quanying Liu, Yue-Jia Luo, Dean Mobbs, Haiyan Wu

## Abstract

Motivated dishonesty is a typical social behavior varying from person to person. Resting-state fMRI (rsfMRI) is capable of identifying unique patterns from functional connectivity (FC) between brain networks. To identify the relevant neural patterns and build an interpretable model to predict dishonesty, we scanned 8-min rsfMRI before an information-passing task. In the task, we employed monetary rewards to induce dishonesty. We applied both connectome-based predictive modeling (CPM) and region-of-interest (ROI) analysis to examine the association between FC and dishonesty. CPM indicated that the stronger FC between fronto-parietal and default mode networks can predict a higher dishonesty rate. The ROIs were set in the regions involving four cognitive processes (self-reference, cognitive control, reward valuation, and moral regulation). The ROI analyses showed that a stronger FC between these regions and the prefrontal cortex can predict a higher dishonesty rate. Our study offers an integrated model to predict dishonesty with rsfMRI, and the results suggest that the frequent motivated dishonest behavior may require a higher engagement of social brain regions.

## 1. Introduction

Dishonesty is defined as the act of deliberately concealing the truth or conveying false information, with the purpose of gaining benefits or avoiding loss (Abe, 2011; DePaulo et al., 2003). Although dishonesty is regarded as a violation of moral code, it is essential for survival and adaptation (Buss, 2019). In the inlab experiments, according to the experimental manipulation, dishonesty can be divided into two types, instructed dishonesty and motivated (or spontaneous) dishonesty (Wu et al., 2009; Sai et al., 2018). For the former, participants make honest or dishonest responses according to the given instructions (Wu et al.,2009); for the latter, participants usually lie spontaneously under material or monetary incentives (Sai et al., 2018). In developmental psychology, motivated lies have appeared in early childhood (Fu & Lee,2007). This kind of dishonesty is driven by multiple motivations. For example, people may commit self-serving dishonesty for their own benefit or prosocial dishonesty for the benefit of others. In this study, we focus on motivated dishonesty driven by self-interest to investigate the individual differences and the neural association underlying it.

Existing task-based fMRI studies have been investigating neural mechanisms of dishonest behavior (either the instructed dishonesty or the motivated dishonesty) and found that dishonesty is associated with many brain areas, including motivation system, cognitive control system, reward system, moral system and so on. For example, dishonesty with selfish motivations leads to increased striatum activity and stronger functional connectivity between valuation and cognitive control network (Pornpattananangkul et al., 2018; Cui et al., 2018), while altruistic dishonesty is considered more acceptable and results in decreased anterior insula (AI) activity (Lewis et al., 2012; Yin et al., 2017). Since dishonesty has to inhibit automatic honest responses or to resist reward temptations, cognitive control is involved in dishonesty (Bargh & Chartrand, 1999; Greene & Paxton, 2009; Haidt, 2001). Evidence from fMRI studies consistently shows that dishonest responses activate brain regions related to cognitive control, such as the dorsolateral prefrontal cortex (DLPFC), the ventrolateral prefrontal cortex (VLPFC), the medial frontal cortex (MFC) and the anterior cingulate cortex (ACC) (Christ et al., 2009; Farah et al., 2014). Moreover, for rewards are usually employed to motivate individuals to lie, the outcomes of successful dishonest choices elicit stronger activation in the reward system, such as the ventral striatum and posterior cingulate cortex (PCC) (Sun et al., 2015). Interestingly, the neural response to the anticipated reward in the nucleus accumbens (Nacc) can significantly predict the dishonest behavior in an incentivized prediction task (Abe & Greene, 2014), indicating that the expectation of reward is critical in motivated dishonesty. In addition to the above processes, morality and self-reference may act as modulators in the middle. For example, participants who paid lower costs for harming others have stronger dorsal striatal responses to the profits gained from doing so (Crockett et al., 2017). Compared with the undetected dishonesty paradigm, the sender-receiver paradigm can arouse stronger self-reference due to the presence of the recipient, leading to increased self-evaluation and neural responses in the orbitofrontal cortex (OFC) and prefrontal cortex (PFC) (Cui et al., 2018; Hughes & Beer, 2013; Yin et al., 2017).

Although task-based fMRI can locate the brain regions under dishonesty, resting-state fMRI (rsfMRI) embodies spontaneous neural activity, and the resting-state brain networks obtained by rsfMRI functional connectivity (FC) can predict individual differences (Biswal et al., 1995; Raichle, 2011). It has been demonstrated that there is a strong overlap between one’s rsfMRI and task-fMRI(i.e., 80%, see (Biswal et al., 1995)). A recent theory proposed that the spontaneous fluctuations of correlated activity within and across brain regions at rest may reflect the most common perceptual, motor, cognitive, and interoceptive states, which are proactive and predictive (Pezzulo et al., 2021). Apart from fundamental cognitive functions (e.g., perception and attention), rsfMRI can also be used to predict complex social behaviors (Li et al., 2013; Tian et al., 2017; Bellucci et al., 2019; Shi et al., 2018; Christov-Moore et al., 2020). One hot topic is whether the pre-task rsfMRI can be used to predict individual dishonesty. Compared with the task-fMRI, rsfMRI is more convenient in data collection, and the protocol is easier if we are interested in predicting lies in daily life (Bettus et al., 2010; Lin et al., 2018). To this end, we will explore whether brain activity in rsfMRI can predict incentive dishonesty. The functional connectivity analysis using rsfMRI has been applied to explore the individual differences in dishonesty, providing neural insights into the cognitive framework of dishonesty. The cognitive processes (including self-reference, cognitive control, reward valuation, and moral regulation) and their interactions are proved to be the main predictors of the frequency of someone’s dishonest behavior (Speer et al., 2020; Yin et al., 2021). Specifically, for brain regions involved in reward processing such as caudate and Nacc, it has been found that their FC with medial PFC (a key region in self-referential processing) is positively correlated with honesty, which suggests the potential of promoting honesty by incorporating internal value into reward evaluation (Yin et al., 2021). This is consistent with the evidence from another task-based fMRI studies, which suggests that the self-processing network, mainly consisting of mPFC, PCC, and temporoparietal junction (TPJ), may maintain a positive self-image and thereby increase honesty (Speer et al., 2020). Moreover, the strengthened FC between the self-referential network and the cognitive control network may prompt honesty (Speer et al., 2020). In addition, the stronger FC between cognitive control network and reward network may be associated with the higher tendency to be dishonest, no direct evidence from rsfMRI studies though (Pornpattananangkul et al., 2018). Moreover, in an integrative framework, it is still unclear how the changes of functional coupling among these four cognitive processes (namely, self-reference, cognitive control, reward valuation, and moral regulation) predict dishonesty. Two rsfMRI-based approaches are commonly used for predicting dishonesty: the connectome-based prediction modeling (CPM) (Ren et al., 2021; Shen et al., 2017) and the priori region-of-interest (ROI) analysis (Yin & Weber, 2019; Yin et al., 2021). Specifically, as a data-driven approach, CPM aims to build the brain-behavior relationship through cross-validated predictive models. Further, the priori ROI analysis defines certain ROIs based on prior knowledge and then establishes the relationship between the FC among ROIs and a specific behavior through multiple comparisons. Despite some rsfMRI studies of (dis)honesty, there is still a lack of comprehensive studies to combine both approaches and compare their results for interpretation. To fill in this gap, we conducted both CPM and a priori ROI analysis to examine to what extent resting-state FC predicts motivated dishonesty.

In this study, we designed an incentive dishonesty task and performed comprehensive rsfMRI analyses to explore the relationship between individual variations in dishonesty behavior and neural patterns, especially the FC between brain regions and networks. In the dishonesty task, we asked participants to send information (food preference) to another player with an assigned reward to motivate the players to pass the true information (honest condition) or to pass the wrong information (dishonest condition). Eight-minute rsfMRI was scanned just before the task. This study focused on the following three aspects: (i) predicting individuals’ dishonest behavior through CPM analysis (**Sec 2.2**); (ii) bridging the individual cognitive processes (self-reference, cognitive control, reward, morality) with dishonesty through ROI analysis and correlation analysis (**Sec 2.3**); (iii) revealing the cooperation and mediation effects existing between these cognitive processes which are associated with dishonesty through the interaction analysis (**Sec 2.4**). Finally, it enables us to form an integrative view on the rsfMRI predictive model for dishonesty based on the findings (See Figure S2 for pipeline of the study).

## 2. Methods

### 2.1. Participants

Thirty-one participants (19 females) participated in the fMRI experiment (age: mean ± *SD* = 24.29 ± 3.81). Two participants were excluded due to excessive head movements (> 2*mm* or > 2°). The remaining 29 participants (17 females) ranged from 18 to 26 years of age (mean ± SD = 22.35 ± 2.03). Participants were recruited from the student community at the university. All participants were righthanded and had no history of neurological or psychiatric disorders. Participants gave written informed consent after being provided with a complete description of the study. Participants received full debriefing on the completion of the experiment. All the procedures involved were in accordance with the Declaration of Helsinki and were approved by the Institutional Review Board (IRB) of the Institute of Psychology, Chinese Academy of Sciences.

### 2.2. Procedure of the task

The task used in the current study was taken from a series of experiments which include seven sessions. From the participants’ perspective, the whole experiment was about information receiving and passing. For this study, we only focused on session 3 of the experiments (an information-passing task). In this task, the participants played the role of information sender, passing the food preference information to another player (receiver) with the consideration of the amount of the reward. Although the participants were told that the receiver might get more reward units if he or she chooses more correct answers based on the received information, there was no real “receiver” to receive the information, and there was no further interaction between the participants and the receiver. In total, we selected 24 pairs of food and randomly chose the preferred food in each pair. The food images were selected from Food-Pics Extended (http://food-pics.sbg.ac.at). Each participant had a learning session to memorize the list of the preferred food. The dishonesty task would not start unless the accuracy of all 24 pairs of food reached above 90%. Thus the participants knew the correct answer for all questions. In each trial (Figure 1a), the question and two answers were presented for 1 s, and then the reward corresponding to each food was presented on the left and right sides. The participants had to choose an answer within 4 s. The chosen answer and the resulting reward would be highlighted for 1 s. The jitters between trials were 2–8 s (mean = 4 s).

**Figure 1:**
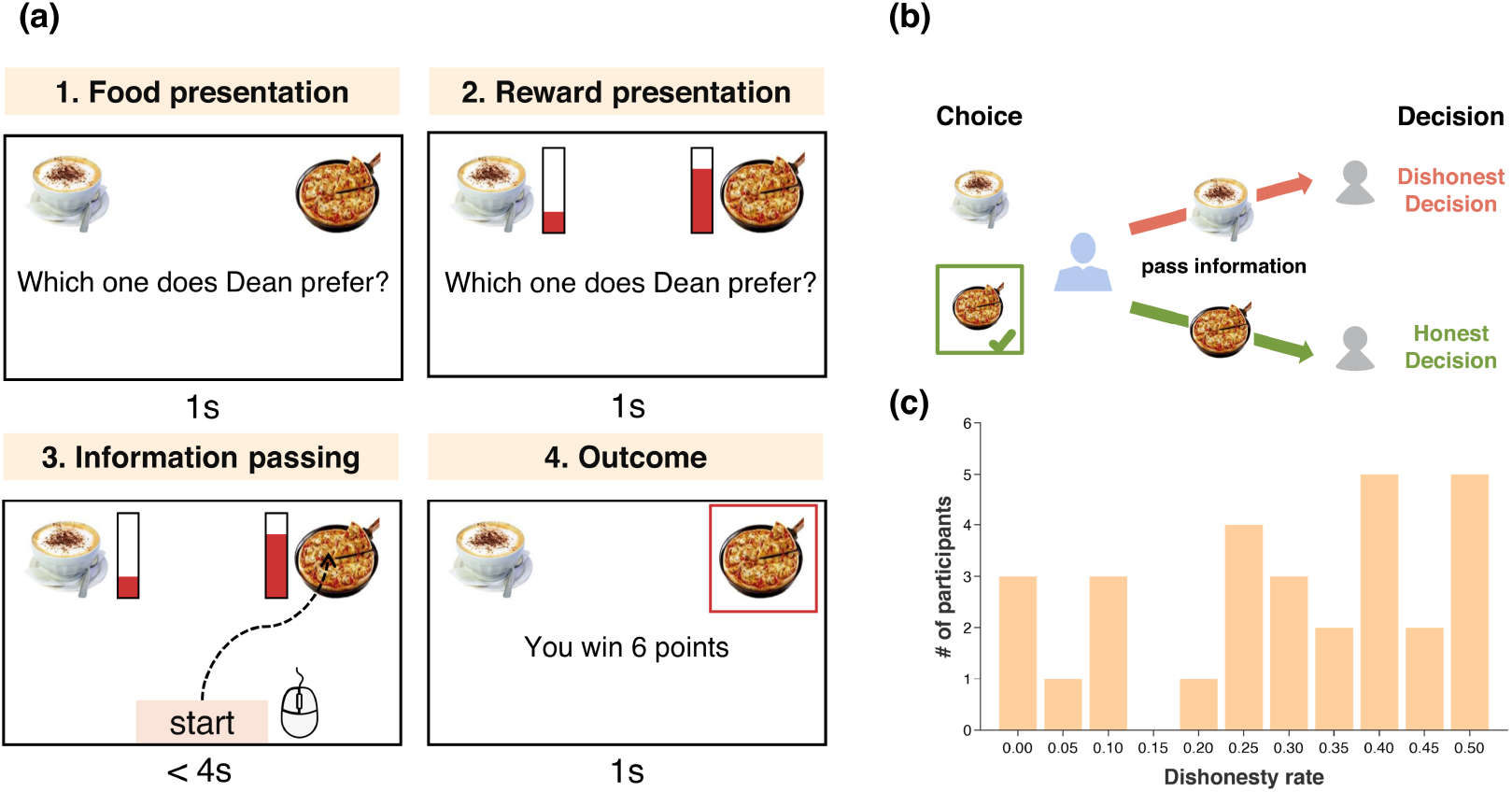
The information-passing task. (a) An example of a single trial. The participant played as an information sender to pass food preference information to another player with the consideration of reward units. (b) Possible responses of participants. Honest decision refers to participants passed the true information based on their memory; Dishonest decision refers to participants passed the false information for more rewards. (c) The histogram of dishonesty rate across participants (*N* = 29).

The trials with a higher reward for the true answer were defined as the **honest condition**. In contrast, the trials with a higher reward for the wrong answer to motivate dishonesty were defined as the **dishonest condition**. The reward units were drawn from a uniform distribution [2, 4, 6, 8], using pseudo-random sampling with replacement. Both the honest and dishonest conditions consisted of 96 trials separated into four runs. Each pair of food items were repeated eight times, with two times in each run. The dishonesty rate was defined as the proportion of false information delivered in the dishonest condition, calculated to measure self-serving lying behavior.

### 2.3. MRI acquisition

MRI data was collected with a General Electric 3T scanner (GE Discovery MR750). The rsfMRI data was collected by an echo-planar imaging (EPI) sequence utilizing gradient echo. Slices were acquired in interleaved order and the data consisted of 200 whole-brain volumes (repetition time (TR) = 2,000 ms, echo time (TE) = 21 ms, flip angle = 90°, slice number = 42, slice thickness = 3.5 mm, matrix size = 64 × 64, field of view (FOV) = 200 mm, and voxel size = 3.1 × 3.1 × 3.5 mm^3^). T1-weighted structural image data was collected for anatomical reference using a 3D magnetization-prepared rapid gradient-echo (MPRAGE) sequence (TR = 2,530 ms, TE = 2.34 ms, flip angle = 7°, FOV = 256 mm, slice number = 176, slice thickness = 1 mm, in-plane matrix resolution = 256 × 256, FOV = 256 mm, and voxel size = 1 × 1 × 1 mm^3^).

### 2.4. fMRI preprocessing

The rsfMRI preprocessing was performed using the advanced DPARSF module V5.1 (Yan et al.,2016). All volume slices were corrected for different signal acquisition times, following each participant’s time series of images. Individual structural images (T1-weighted MPRAGE) were co-registered to the mean functional image after realignment. The transformed structural images were segmented into the gray matter (GM), the white matter (WM), and cerebrospinal fluid (CSF) (Ashburner & Friston, 2005). To remove the nuisance signals, the Friston 24-parameter model (Friston et al., 1996) was utilized to regress out head motion, as well as the mean WM and CSF signals. The functional data from individual native space was transformed to the standard Montreal Neurological Institute (MNI) space (http://www.bic.mni.mcgill.ca/ServicesAtlases/HomePage). Spatial smoothing (FWMH kernel: 6 mm) was applied to the functional images for the connectivity analysis. Further, temporal filtering (0.01–0.1 Hz) was performed on the time series.

### 2.5. CPM: Support Vector Regression

The functional connectivity matrix construction was conducted in the DPARSF toolbox. First, we extracted 264 ROIs in the whole brain according to the Power atlas (Power et al., 2011). 264-by-264 functional connectivity (FC) matrix was obtained by Pearson’s correlation analysis between the averaged BOLD time series of each pair of ROIs. Fisher’s Z transform was performed on Pearson’s correlation coefficient (*r*). The static FC matrix was calculated participant by participant, resulting in 29 FC matrices. Then, a support vector regression (SVR) analysis was applied to the static FC matrices to predict the individual dishonesty rate using leave-one-out cross-validation (LOOCV) (Beaty et al., 2018;Dosenbach et al., 2010; Finn et al., 2015). (10-fold cross-validation was used to verify the results, see supplementary materials for more details). For each iteration of LOOCV, participants were divided into training sets with 28 samples (*N*-1) and test sets with one sample. Feature selection was applied only to the training set to obtain relatively sparse features. The criterion for feature selection was that the edges should be significantly correlated to dishonesty rate with *p* < 0.01 (Rosenberg et al., 2016; Takagi et al.,2018). All features of 28 participants (N – 1) were trained using SVR, and the features of the left-out participant were tested on the trained model to output a predicted value of dishonesty rate. LOOCV was performed 29 times (N), with each participant being the test sample once. Then the 29 predicted dishonesty rates were correlated with the actual dishonesty rates, and the Spearman correlation *ρ* and p-value along with mean squared error (MSE) were used to evaluate model performance (Figure 2).

**Figure 2:**
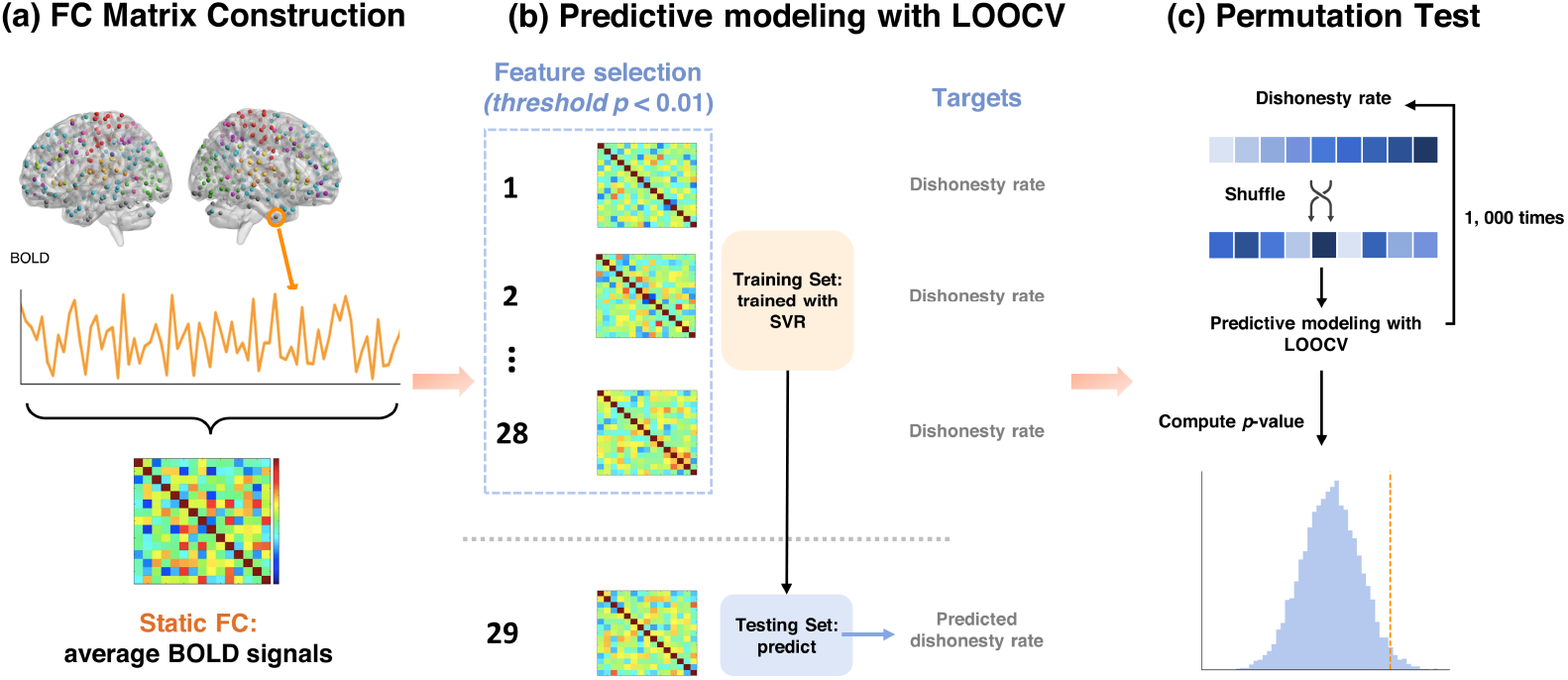
Workflow for Connectome-based Predictive Modeling. (a) FC matrix construction. The time series of each ROI defined in the Power atlas was extracted the averaged (Power et al., 2011). Pearson correlation was used to compute the FC matrix between each pair of ROIs. (b) Predictive modeling with LOOCV. Specifically, SVR was used to train the FC matrices from *N* – 1 participants; the trained SVR model was tested on the left-out one participant’s FC matrix and output a predicted dishonesty rate. (c) Permutation test for statistics. The significance of prediction was evaluated by conducting 1,000 permutations.

264 ROIs in the Power atlas can be grouped into 13 functional networks, namely somatomotor network (SOM), auditory network (AUD), visual network (VIS), default mode network (DMN), memory retrieval network (MRN), dorsal attention network (DAN), ventral attention network (VAN), salience network (SN), cingulo-opercular task control network (COTCN), fronto-parietal task control network (FPTCN), subcortical network (SUB), cerebellar network (CER) and the uncertain regions. To quantify the predictive power between and within networks, we summed the absolute value of regression weights for functional connectivity between the same network pair. The results were visualized in a 13-by-13 matrix, with each entry denoting a network pair.

The permutation test was conducted to evaluate the statistical significance of the predictive model. Specifically, the actual dishonesty rates from 29 participants were randomly shuffled 1,000 times. For each time of permutation, the shuffled model was trained in 28 participants and then output a prediction of the dishonesty rate of the remaining one participant. This procedure was repeated 29 times, resulting in 29 predicted dishonest rates. Then we calculated the correlation coefficient (*ρ*) between the predicted and actual dishonesty rates. Finally, we counted the number of *ρ* values from the 1,000 permutations greater than the real ones from the true model and divided it by the times of permutation (1000), obtaining the *p* value of the permutation test. To further examine the generalization of the present model (Poldrack et al., 2020; Scheinost et al., 2019; Zuo et al., 2019), we have also performed the external validation with another fMRI data of dishonesty task (see supplementary materials for more details).

### 2.6. ROI Analysis

Based on the cognitive processes involved in dishonesty and previous studies using task-related fMRI and rsfMRI to reveal the neural basis of dishonesty, ROIs were chosen from self-reference, cognitive control, reward, and morality, four distinct social brain network. ROIs in the four networks were generated by the meta-analyses on Neurosynth (https://neurosynth.org/). For example, the ROI in the self-referential system was obtained by a 10-mm spherical mask centered in the peak coordinate from the meta-analysis term “self-referential” in Neurosynth. The ROI analysis was implemented in AFNI (http://afni.nimh.nih.gov/). The mean time course of all voxels in each ROI was extracted and correlated with the time course of every voxel in the brain. The correlation between time courses was transformed to Fisher’s Z-score and used as FC. Then, the FC between ROI and each voxel was correlated with the dishonesty rate. There were 47355 voxels in the whole brain mask, correlations have been tested for every ROI-voxel pair for each ROI. Voxel-wise threshold was set to uncorrected p < 0.001as recommended and the cluster-wise threshold was set to multiple comparison corrected p < 0.01 at group level (Cox et al.,2017; Yin et al., 2021). To further control false-positive rates, a threshold for cluster size was used (Cox et al., 2017). The threshold was calculated according to the voxel-wise and cluster-wise threshold, in the current study was 64 voxels.

### 2.7. Network Interaction Analysis

We conducted network interaction analysis to investigate how social brain networks interacted directly and whether the FC between networks could predict dishonesty. Moreover, the interaction analysis enabled us to test the potential mediation effects underlying FC between networks and dishonesty rate. Masks of self-referential, cognitive control, reward, and moral network were obtained from meta-analyses on Neurosynth, expending to the whole network. xjView (https://www.alivelearn.net/xjview) was used to generate new masks with threshold Z > 5 to ensure the stability of each brain network. Time series of all voxels in the mask were extracted and averaged. Then FC was calculated between each pair of networks and converted to Fisher’s Z-score using DPARSF. Finally, the relationship between Fisher’s Z-score and the dishonesty rate was obtained by Pearson’s correlation. Considering the overlap between different brain networks, we conducted conjunction analysis to ensure that the effects were not caused or significantly influenced by the overlap. The overlapped brain region between each pair of networks was calculated and subtracted from the corresponding network mask. The previous interaction analyses were repeated based on the subtracted masks to eliminate the impact of the overlapping.

## 3. Results

### 3.1. Behavioral Analysis

The overall dishonesty rate for the two conditions in the information-passing task of 29 participants ranged from 0.004 to 0.51 (mean ± *SD* = 0.30 ±0.16). The distribution of the dishonesty rate of all participants is shown in Figure 1c, manifesting pronounced individual differences. For debrief questions, all participants declared they did not doubt the cover story and did not know the true purpose of the experiment.

### 3.2. CPM analysis

CPM was used to examine the degree to which the resting-state FC could predict dishonesty. Correlation matrices for static FC were built respectively, and support vector regression (SVR) was selected for training static FC. Cross-validated models demonstrated that static FC can significantly predict the dishonesty rate of participants (Figure 3a; *MSE* = 0.023, *ρ* = 0.421, p = 0.023). The permutation test was conducted to examine the significance of the result (Figure 3b), and it demonstrated that the correlation for static FC is significantly greater than the chance level (p = 0.016). Using features from the internal validation model, the external validation led to a significant true model and a significant predictive effect in the permutation test (Figure S4).

**Figure 3:**
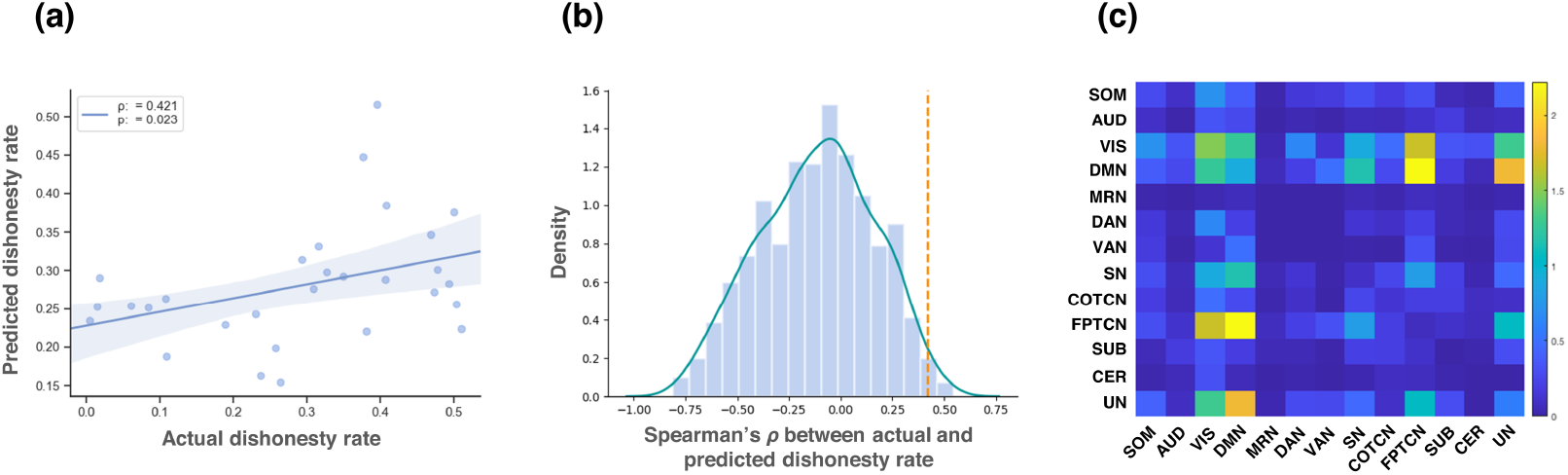
Performance of CPM on dishonesty prediction. (a) The Spearman’s *ρ* between the actual dishonesty rate and the predicted dishonesty from the CPM model. (b) The histogram of the Spearman’s *ρ* values after 1,000 permutations. The dashed orange line indicates Spearman’s *ρ* of the true model. The permutation test resulting *p* value is 0.016. (c) The absolute values of predicting coefficient between and within the same pair of networks were summed and visualized in a 13-by-13 coefficient matrix according to network partition of Power atlas. *Abbrev*.: SOM, somatosensory network; AUD, auditory network; VIS, visual network; DMN, default mode network; MRN, memory retrieval network; DAN, dorsal attention network; VAN, ventral attention network; SN, salience network; COTCN, cingulo-opercular task control network; FPTCN, fronto-parietal task control network; SUB, subcortical network; CER, cerebellum network; UN, uncertain.

Moreover, to calculate the magnitude of network-wise coefficient in predictive modeling, the absolute coefficients for edges connecting the same pair of networks were summed. The result showed that the sum coefficient is highest for edges connecting DMN and FPTCN (Figure 3c). Comparisons between the coefficient matrix of CPM

### 3.3. ROI Analysis

ROI of each network was selected from the most strongly activated cluster of meta-analysis. For the self-referential network, individuals with higher dishonesty rate showed increased FC between ROI (x = 0, y = 58, z = 16) and left superior orbital gyrus (Figure 4a; peak MNI coordinate: 0, 51, 24; 220 voxels), right middle frontal gyrus (peak MNI coordinate: 30, 48, 3; 165 voxels), and left inferior frontal gyrus (peak MNI coordinate: 48, 21, −3; 73 voxels). For the cognitive control network, individuals with higher dishonesty rate showed increased FC between ROI (x = 4, y = −26, z = 30) and left superior medial gyrus (Figure 4b; peak MNI coordinate: 3, 45, 24; 72 voxels). DMN and FPTCN contributed to the processing of self-related information and cognitive control, respectively. In terms of large-scale brain networks, the results from CPM demonstrated that the enhanced functional coupling between DMN and FPTCN can predict the dishonesty rate. In terms of specific social brain networks, the results of ROI analysis revealed that FC between the frontal cortex and ROIs are correlated with dishonesty rate. Both results proved that brain networks indirectly or directly involved in self-referential and cognitive control are the basis of dishonest behavior.

Furthermore, in order to explore the possibility of other cognitive processing involved in dishonesty, ROIs were extracted from reward network and moral network on Neurosynth. For the reward ROI (x = 10, y = 8, z = −8), its FC with left superior medial gyrus (Figure 4c; peak MNI coordinate: −3, 54, 39; 144 voxels) was positively correlated with dishonesty. And for the moral ROI (x = 4, y = 62, z = 14), such positive correlation with dishonesty was found in the FC between moral ROI and left superior frontal gyrus (Figure 4d; peak MNI coordinate: −18, 57, 6; 239 voxels), and right middle frontal gyrus (peak MNI coordinate: 30, 48, 3; 139 voxels).

**Figure 4:**
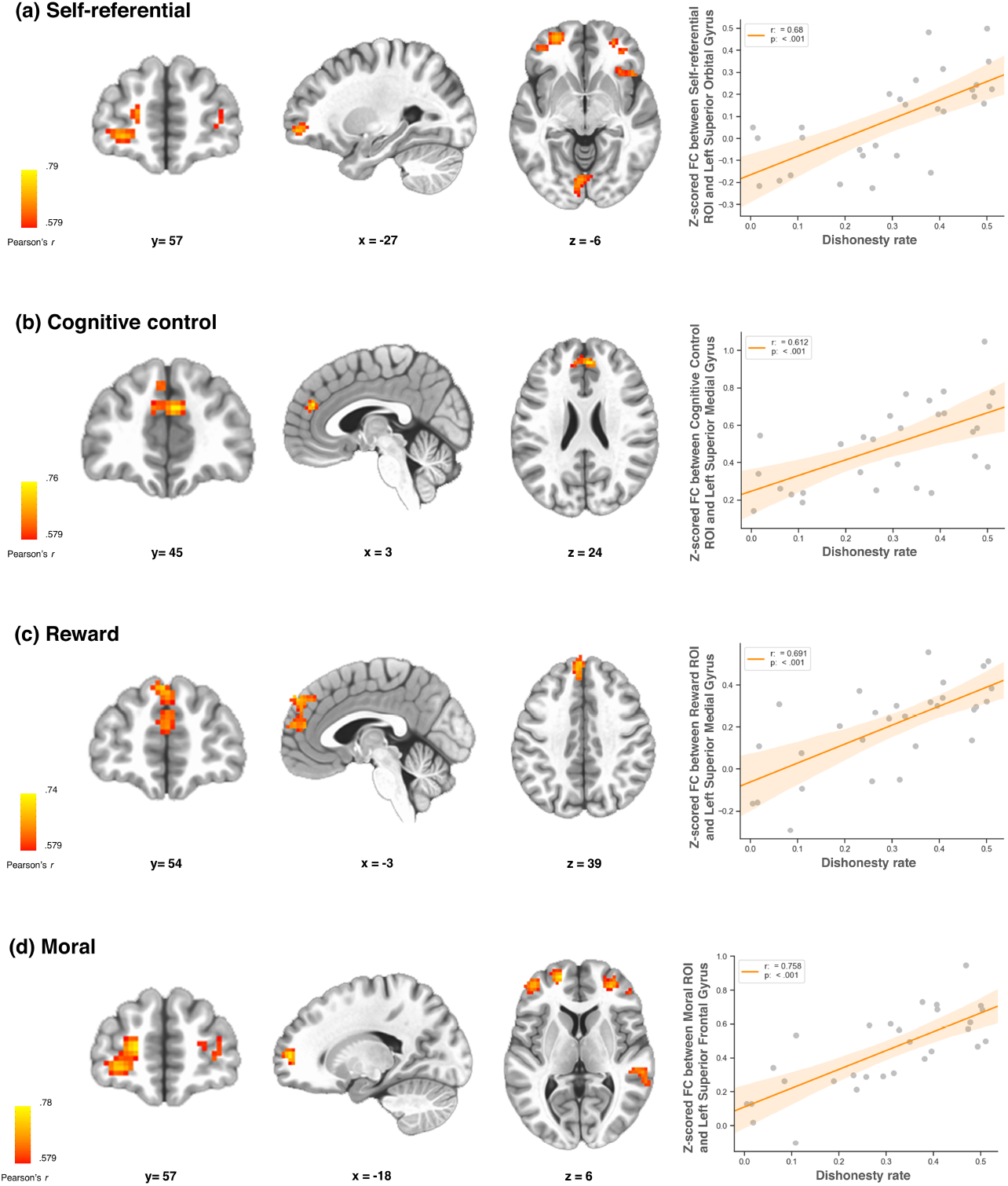
Functional connectivity between PFC and the clustered regions correlated with dishonesty. (a) The FC between self-referential ROI (x = 0, y = 58, z = 16) and left superior orbital gyrus was positively correlated with dishonesty rate (*r* = 0.680, *p* < 0.001). (b) The FC between cognitive control ROI (x = 4, y = −26, z = 30) and left superior medial gyrus was positively correlated with dishonesty rate (r = 0.612, p < 0.001). (c) The FC between reward ROI (x = 10, y = 8, z = −8) and left superior medial gyrus was positively correlated with dishonesty rate (r = 0.691, p < 0.001). (d) The FC between moral ROI (x = 4, y = 62, z = 14) and left superior frontal gyrus was positively correlated with dishonesty rate (r = 0.758, p < 0.001). The detailed information of all clusters associated with each ROI is listed in Table 1.

**Table 1:** Functional connectivity which is significantly correlated with dishonesty rate.

|  |  |  |  |  |  |
| --- | --- | --- | --- | --- | --- |
| **ROI regions** | **MNI coordinates** |  |  |  | **p-value** |
| **Cluster peak** | **voxels** | **x** | **y** | **z** |  |
| **Self-referential (spherical mask)** |  |  |  |  |  |
| Left superior orbital gyrus | 220 | -27 | 57 | -6 | < 0.001 |
| Right middle frontal gyrus | 165 | 30 | 48 | 3 | < 0.001 |
| Left inferior frontal gyrus | 73 | 48 | 21 | -3 | < 0.001 |
| **Cognitive control (spherical mask)** |  |  |  |  |  |
| Left superior medial gyrus | 72 | 3 | 45 | 24 | < 0.001 |
| **Reward (spherical mask)** |  |  |  |  |  |
| Left superior medial gyrus | 144 | -3 | 54 | 39 | < 0.001 |
| **Moral (spherical mask)** |  |  |  |  |  |
| Left superior frontal gyrus | 239 | -18 | 57 | 6 | < 0.001 |
| Right middle frontal gyrus | 139 | 30 | 48 | 3 | < 0.001 |
| Right middle temporal gyrus | 98 | 63 | -36 | 3 | < 0.001 |
| Right insula lobe | 83 | 33 | 21 | -3 | < 0.001 |
| Right infeior frontal gyrus | 74 | 54 | 21 | 30 | < 0.001 |
| Left superior medial gyrus | 65 | -3 | 21 | 42 | < 0.001 |
| Note: voxel size is $3 \times 3 \times 3mm^3$ | | | | | |

### 3.4. Network interaction Analysis

Network interaction analyses were applied to explore how different brain networks interact in predicting dishonesty rate under resting-state. The results showed that FC between self-referential network and cognitive control, reward, moral network were positively correlated with dishonesty rate. The same trend was found in FC between reward and moral network with dishonesty rate (*p* < 0.05, uncorrected; Figure 5a). The coupling between networks which had the highest correlation coefficient with dishonesty rate were plotted in Figure 5b (self-referential and cognitive control network, *r* = 0.491, p = 0.007) and 5c (self-referential and moral network, *r* = 0.509, p = 0.005). To eliminate the influence of conjunction, we further explored if these effects are still significant after controlling the common region of each pair of brain networks. The effect size and direction did remain the same in the conjunction analyses (Figure S6).

**Figure 5:**
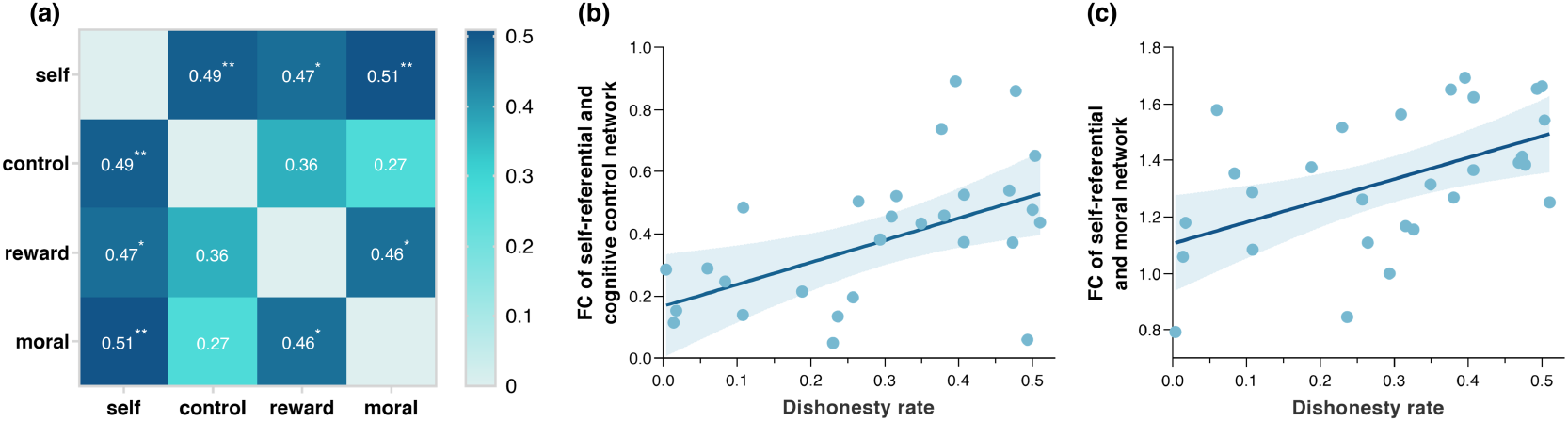
Functional connectivity between the four networks correlated with dishonesty. (a) Heat map for correlation analysis between brain network FC and dishonesty rate. The correlation coefficients were listed in each cell with significance level: * represents *p* < 0.05, ** represents *p* < 0.01. (b) The FC between self-referential network and cognitive control network was positively correlated with dishonesty rate. (c) The FC between self-referential network and moral network was positively correlated with dishonesty rate.

## 4. Discussion

The present study explored whether rsFC can predict individuals’ social decisions like motivated dishonesty. We collected rsfMRI and asked participants to perform an information passing task with monetary reward temptation. Through estimating prediction from individuals’ intrinsic fMRI FC to spontaneous lying, we identified that the rsFC can predict motivated dishonesty from many aspects of correlations. The current work provides further evidence linking individuals’ moral decisions and rsFC in key networks based on previous findings (Greene & Paxton, 2009; Yin & Weber, 2019; Yin et al.,2021). Moreover, we found distinct patterns of association between rsFC in different brain networks and individuals’ choices in social decisions.

In the whole-brain analysis, results of CPM suggest the FC between DMN and FPTCN can positively predict dishonesty. Several lines of evidence can support this link. We speculate that the connection to fronto-parietal regions is related to executive function: including conflict monitoring and resolution(Marek & Dosenbach, 2018; Wang et al., 2010; Jiang et al., 2018; Brydges et al., 2020). Also, it is found that fronto-parietal regions may play a role in rationalization, which can resolve cognitive dissonance (Jarcho et al., 2011). This is consistent with the notion that lying requires recruitment of cognitive resources for conflict resolution and rationalization (Van’t Veer et al., 2014), so frequent lying may result in a greater likelihood of exerting top-down control on other related cognitive networks. DMN has been identified by previous studies as a functional network to generate internally-direct thought and process self-related information (Molnar-Szakacs & Uddin, 2013; Qin & Northoff, 2011), so the strengthened FC between FPTCN and DMN may be associated with more dishonesty by resolving the moral conflict, rationalizing the dishonest act, and internalizing the rationalization. Therefore, the enhanced cooperation between cognitive control and self-related processes may help people better adapt to lying.

Our findings of the ROI analysis can complement the CPM results, providing a more comprehensive explanation for the cognitive framework for dishonesty in the resting state. It confirms the positive correlation between multiple rsFC networks and dishonesty. For example, the four cognitive networks may be associated dishonesty through FC with specific regions in PFC. Specifically, for the self-referential network (Figure 4a), its FC with OFC and DLPFC was positively correlated with dishonesty. Since it is reported that DLPFC exerts top-down control(Milham et al., 2003; Brosnan & Wiegand, 2017; Prutean et al., 2021), we interpret this enhanced FC as top-down control imposes on the self-related processes. Similarly, for cognitive control network (Figure 4b), the increased FC with mPFC, a core region in self-referential processing(Lieberman et al., 2019; D’Argembeau et al., 2007), predicts a higher dishonesty rate. The enhanced FC between self-referential and cognitive control processes found in both CPM and ROI analysis may reflect the importance of this cooperative relationship in dishonesty. As two key cognitive processes, self-reference and cognitive control have been examined separately in previous dishonesty studies (Yin et al., 2021; Ding et al., 2017; Gombos, 2006). For instance, self-referential processing may be associated with increased honesty by maintaining a positive self-image (Yin et al., 2021), while the recruitment of cognitive control network is found in both honest and dishonest behaviors(Ding et al., 2017;Gombos, 2006; Johnson Jr et al., 2004; Walczyk et al., 2003). A recent study has provided reconciling evidence that the influence of cognitive control in (dis)honesty may vary depending on one’s moral default, and cognitive control is recruited to override moral default (Speer et al., 2020). Our findings further add to the evidence of the recruitment of cognitive control in dishonesty. In particular, the cognitive control process may be recruited to cooperate with self-referential processing to internalize rationalized content and promote dishonesty (Jarcho et al., 2011).

Moreover, ROI analysis on reward network shows that increased FC with mPFC is associated with dishonesty (Figure 4c). Since mPFC is involved in value representation(Gross et al., 2014; Wang et al.,2014; Gazit et al., 2020), the increased FC between mPFC and reward network may imply the encoding of a higher reward value, which will motivate dishonest behavior. ROI analysis of the moral network revealed that the FC with regions in DLPFC, temporal lobe, insula, and inferior frontal gyrus (IFG) are associated with dishonesty rate (Figure 4d). This large-scale network may suggest a more complicated relationship between morality and dishonesty. In predicting dishonesty, the largest cluster functionally connected to the moral network was found in DLPFC. DLPFC can implement flexible goal types, promoting either selfish (FeldmanHall et al., 2012; Yamagishi et al., 2016)or moral (Knoch et al., 2009; Strang et al.,2015)goals across different contexts. So for the paradigm in which participants can gain monetary rewards by lying, we posit that DLPFC may prioritize pursuing self-interest and thus exert control on the moral network.

Interaction analyses show that the FC between the self-referential network and the other three networks and the FC between reward and moral network can positively predict dishonesty rate (Figure 5a). These findings suggest that mutual cooperation between these cognitive processes may increase dishonesty. We speculate that the self-referential network is not only directly associated with dishonesty but also bridges the neural patterns of other networks. To test this hypothesis, we further conducted a mediation analysis and found that FC between self-referential and moral networks can completely mediate the predictive effect of the FC between reward and moral on dishonesty rate (Figure S7). This mediation may imply that when making dishonest decisions, processing of self-related information may interfere with the computation of moral value (Diener & Wallbom, 1976; Ruff & Fehr, 2014; Yin et al., 2021). Combined with ROI-based analysis, we infer that the superior medial gyrus which resides in mPFC might act as a hub to connect networks and brain regions involved in dishonesty, including caudate, dlPFC, PCC, and insula. However, our results are different from the previous study which found the inferior frontal gyrus as a potential hub to facilitate dishonest responses (Yin & Weber, 2019). The different paradigms may explain this difference in the two studies. Their task was focused on self-serving dishonesty(Yin et al., 2017); in contrast, we adopted the information passing task and used rewards to induce dishonest behaviors of participants, so the subjects needed to be more self-disciplined (Cui et al., 2018; Pornpattananangkul et al., 2018).

There are several limitations in the current study. First, the behavioral task used to measure personal dishonesty in this study (Figure 1) is more complex than common paradigms. Participants can not lie in private but have to deceive others for more rewards, which may force them to pay more attention to self-image and reputation (Cui et al., 2018). Besides, excessive image stimulus in the task may have a certain impact on the judgment, which can partially explain the confirmed effects of the visual network in the CPM analysis. Further, the FC is measured by Pearson correlation which cannot reflect the directionality of the interactions between brain regions and networks. SVR or other regression models cannot reveal the causal effects of FC on dishonesty. Therefore, the modulatory role of certain brain regions and systems cannot be determined. For example, we found that the FC between cognitive control and reward valuation network can positively predict dishonesty rate. However, without examining the directionality, we could not determine whether the relationship is bottom-up or top-down. Most importantly, although the features of CPM analysis survived the permutation test and external validation (see supplementary materials for details), the relatively small sample size not only led to a small effect size but brought bias and variance to the model (Scheinost et al., 2019). Besides, according to a recent preprint, a sample size of about 200 is required for the reproducibility of models between rsFC and cognitive ability (Marek et al., 2020). Furthermore, this study only focused on the cognitive system of dishonesty while ignoring the emotion system. According to previous studies, individuals cannot avoid emotion fluctuations in the process of lying, therefore emotion can be an important indicator of lie detection (Greene & Haidt, 2002;Moll et al., 2002). How the cognition and emotion systems interact in the process of lying and whether these interactions can be manifested at the neural level remains to be explored in the future.

## 5. Conclusion

In summary, both CPM and ROI analysis demonstrated that FC during resting-state can predict dishonesty. In CPM, the increased FC between DMN and FPTCN can strongly predict dishonesty, indicating top-down control of self-related processing. In ROI analysis, the four cognitive processes (self-reference, cognitive control, reward valuation, and moral regulation) can predict dishonesty through the FC with specific regions in PFC such as dlPFC, mPFC, and IFG, respectively. The results reflect an important role of PFC in promoting dishonest behavior. Further analyses showed that there might be cooperation and mediation between these cognitive processes that lead to dishonest behaviors.

## Supporting information

see supplementary materials for more details

## Data and Code Availability Statement

The data used in this manuscript is not available due to privacy issues. The code used in this manuscript is available at https://github.com/andlab-um/restDishonesty.

## Acknowledgement

This study is surpported by SRG of UM:SRG2020-00027-ICI, National Natural Science Foundation of China: U1736125 and Natural Science Foundation of Guangdong Province (2021A1515012509, 2019A1515111038). The authors would like to thank Mr. Hao Yu who provided general support in participants recruiting.

## Conflict of interest

All authors declare no competing interests.

## Notes

### Competing Interest Statement

The authors have declared no competing interest.

