## Supplementary material for "Resting-state functional connectivity of social brain regions predicts motivated dishonesty": see supplementary materials for more details


1. **Brief introduction of the whole study**
2. **Pipeline of the current study**
3. **Results of 10-fold internal validation**
4. **External validation**
5. **Comparison between the coefficient matrix of CPM**
6. **Conjunction analysis of brain networks**
7. **Mediation analysis of self-referential network**


### 1. <u>Brief introduction of the whole study</u>

The whole study is a within-subject design, with seven sessions (six sessions on the fMRI scanning day while one session for three days after the fMRI scanning) in total. The paradigm of the whole study is shown in Figure S1.

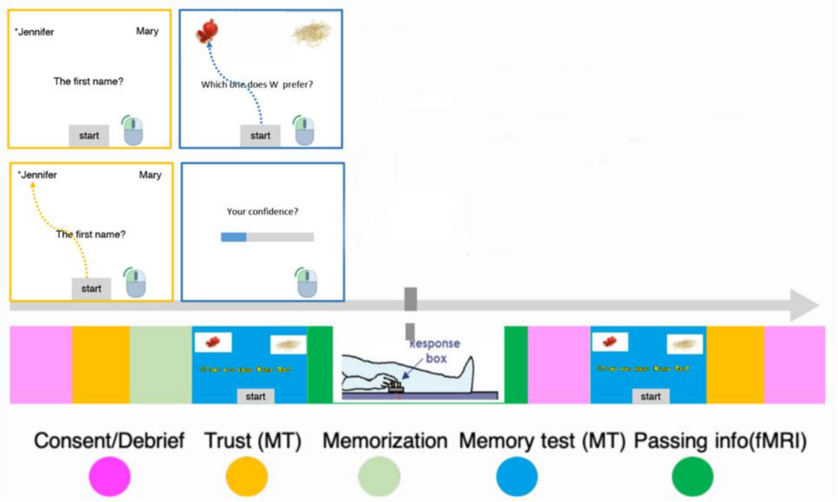


**Figure S1** *The paradigm of the whole study.* MT means mouse tracking.

**Session 1:** Information receiving and trust decisions. Participants receive information from an anonymous player who has memorized the information and decided whether to trust the information. Note that the information is different from the information of the subsequent sessions. This session aims to make the participants believe in the protocol (receive, memorize, and pass information) and obtain a measurement of baseline trust level.

**Session 2:** Food preference memory test. Participants need to memorize someone's food preference and reach 100% accuracy in the test. This session is a preparation for session 3.

**Session 3:** Information-passing task in the scanner (see detailed description in the main text), which is the main task we try to build the predictive model.

**Session 4:** "Exit questionnaire". Participants finish questionnaires and some debrief questions. The purpose of this session is to acquire an explicit measure of the motives and make the natural "forgetting" happen by telling participants the experiment is almost finished.

**Session 5:** Surprise second memory test. Participants take the second memory test surprisingly. The purpose of this session is to test the memory change after the dishonesty task.

**Session 6**: Second information receiving. The procedure and instruction are the same as session 1. The purpose of this session is to measure the possible trust change effect after the honest/dishonest responses in session 3.

**Session 7** (three days post the scanning): Third memory test. The procedure and instruction are the same as in session 5. The purpose of this session is to test the lasting effect of memory change after the dishonesty task.

### 2. Pipeline of the current study

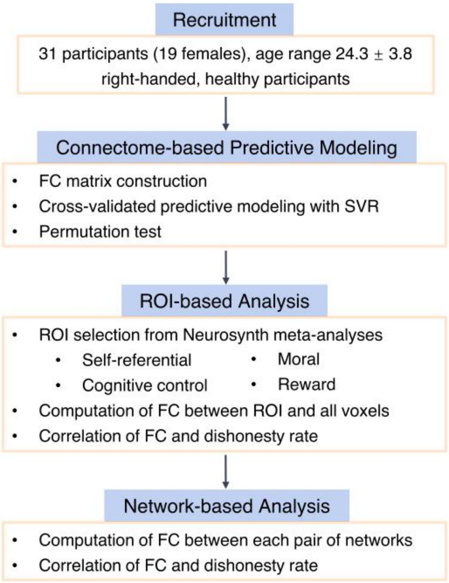


**Figure S2** *Pipeline of the current study.* Analyses of rsfMRI functional connectivity were conducted from three aspects, including connectome-based predictive modeling, ROI-based analysis, and network-based analysis.

### 3. Results of 10-fold internal validation

10 times 10-fold cross-validation was used to verify the results. For each iteration of 10-fold, participants were divided into training sets with 26 or 27 samples (9/10 of the sample size) and test sets with 2 or 3 samples (1/10 of the sample size). To avoid the randomness of once division, 10-fold was repeated 10 times. Then, feature selection, modeling, and evaluation were applied following the same procedure as leave-one-out cross-validation in the main text. 10-fold cross-validated model demonstrated that static FC can significantly predict the dishonesty rate of participants (Figure S3a, $MSE = 0.027$, $\rho = 0.30$, $p < 0.001$). The permutation test demonstrated that the model performance is significantly greater than the chance level (Figure S3b, $p = 0.04$).

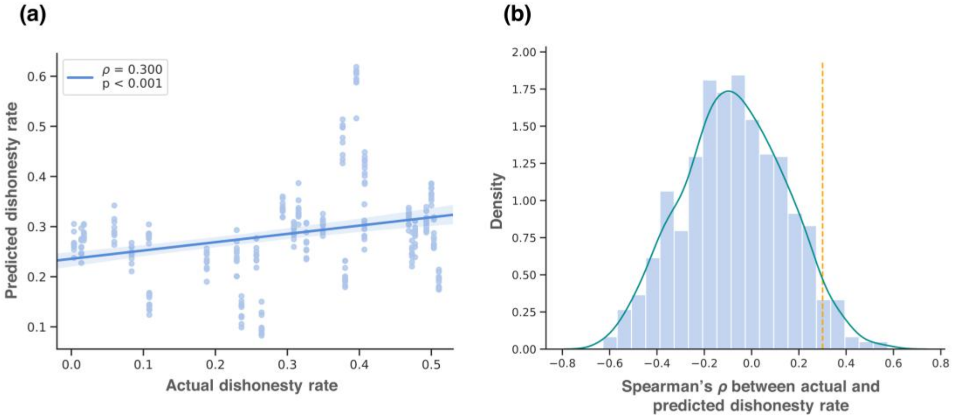


**Figure S3** *Results of 10-fold internal validation.* (a) The Spearman's $\rho$ between the actual dishonesty rate and the predicted dishonesty from the CPM model. (b) The histogram of the Spearman's $\rho$ values after 1,000 permutations. The dashed orange line indicates Spearman's $\rho$ of the true model. The permutation test resulting $p$ value is 0.04.

### 4. External validation

#### Participants

29 healthy participants (14 female; age*: mean* ± *SD = 19.86* ± *3.69*) from the University of Macau were recruited by online advertisement. One participant was excluded due to excessive head movements (>2mm or >2°). Three participants were excluded because they remained honest throughout the whole experiment. All participants were right-handed with normal or corrected-to-normal vision and had not participated in any similar studies. Each participant signed informed consent before the formal experiment, and the experimental protocol was approved by

the Institutional Review Board of the University of Macau (BSERE21-APP005-ICI). At the end of the study, the participants were paid a 130 - 150 mop supermarket coupon.

**Procedure of the task**

The experiment was a self-paced information-passing task with nine blocks. Each block consisted of the same set of 20 trivia questions (e.g., What is the main component of the Martian atmosphere? ) in randomized order. Participants were instructed to pass an answer to another player considering the reward units and his or her past performance. In each trial, correct and incorrect answers were shown separately on the corner of the screen. The correct answer was marked by a black circle. Apart from the correct answer, the monetary reward of each choice and how many times participants had chosen the choice in the former blocks were also provided. To induce participants to make a trade-off between money and honesty, a higher reward was set for the incorrect answer in over 60% of trials. Participants had 4 s to answer, feedback was shown after their choice and lasted for 1 s. If participants failed to make a valid response within 4 s, warnings would display, and that trial would be abandoned in the further analysis. After participants finished each block, the cumulative monetary reward in this block would be displayed on the screen. Same as the main task we used in the main text, the trials with a higher reward for the wrong answer to motivate dishonesty were defined as the dishonest condition. The dishonesty rate was defined as the proportion of false answers delivered in the dishonest condition, calculated to measure self-serving dishonest behavior.

**MRI acquisition**

All MRI data were acquired using a 3.0 T Siemens MAGNETOM Prisma MRI scanner with a 64-channel head coil at the Center for Cognitive and Brain Sciences at the University of Macau. High-resolution T1-weighted images were acquired for each participant before at first (3D MPRAGE sequence; voxel size = 1 mm × 1 mm × 1 mm; FOV = 256 mm; 176 slices, slice thickness: 1 mm; TR = 2300 ms, TE = 2.26 ms, flip angle = 8°). An eight-minute resting-state fMRI was collected by an echo-planar imaging (EPI) sequence before the task (voxel size = 2 mm × 2 mm × 2 mm; FOV = 256 mm; 63 slices, slice thickness: 2 mm; TR = 1000 ms, TE = 30 ms, flip angle = 90°).

**fMRI preprocessing**

fMRI preprocessing remained the same as the main text.

**CPM method**

CPM method remained the same as the main text except that leave-one-out cross-validation was replaced by 10-fold cross-validation. For each iteration of 10-fold, participants were divided into training sets with 22 or 23 samples (9/10 of the sample size) and test sets with 2 or 3 samples (1/10 of the sample size). To avoid the randomness of one division, 10-fold was repeated 10 times.

**Results**

10 times 10-fold cross-validated model demonstrated that static FC can significantly predict the dishonesty rate of participants (Figure S4a, *MSE* = 0.016, $\rho = 0.34$, $p < 0.001$). The permutation test was conducted to examine the significance of the result, and it demonstrated that the model performance is significantly greater than the chance level (Figure S4b, $p = 0.03$).

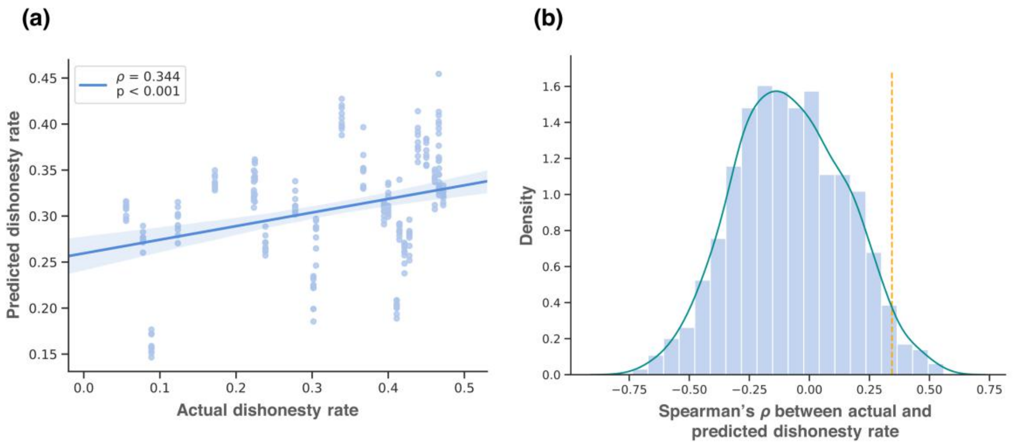


**Figure S4** *Results of 10-fold external validation.* (a) The Spearman's $\rho$ between the actual dishonesty rate and the predicted dishonesty from the CPM model. (b) The histogram of the Spearman's $\rho$ values after 1,000 permutations. The dashed orange line indicates Spearman's $\rho$ of the true model. The permutation test resulting $p$ value is 0.03.

### 5. Comparison between the coefficient matrix of CPM

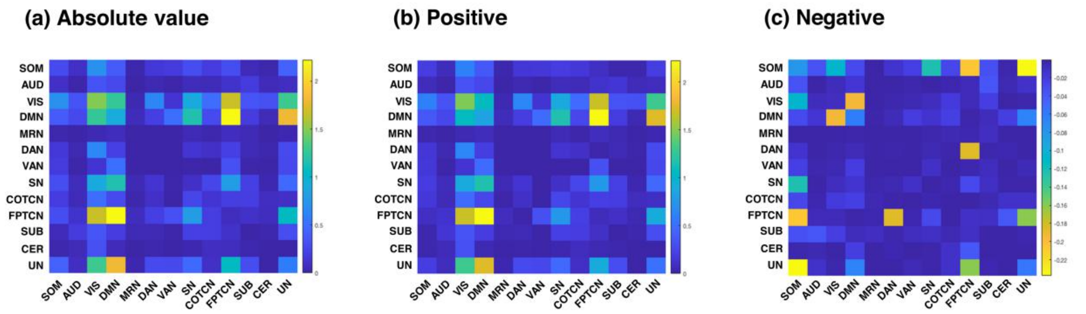


**Figure S5** *Comparison between the coefficient matrix of CPM.* In the main text, absolute coefficients were summed to visualize the predictive power between and within networks. The positive coefficient matrix plotted by the same method was consistent with the absolute coefficient matrix. Besides, the negative coefficient matrix had a small effect size, indicating that the direction of prediction did not significantly impact the result of CPM.

### 6. Conjunction analysis of brain networks

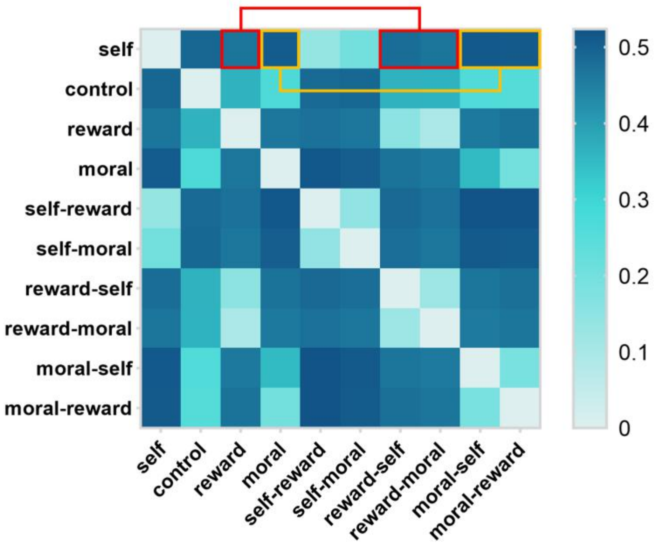


**Figure S6** *Conjunction analysis of brain networks.* To ensure that the interaction effects were not caused or significantly influenced by the overlap between networks, the common region was calculated and subtracted from the corresponding mask. The results of repeated interaction analysis remained the same in both effect size and direction, as highlighted by the red and orange boxes in the figure.

### 7. Mediation analysis of self-referential network

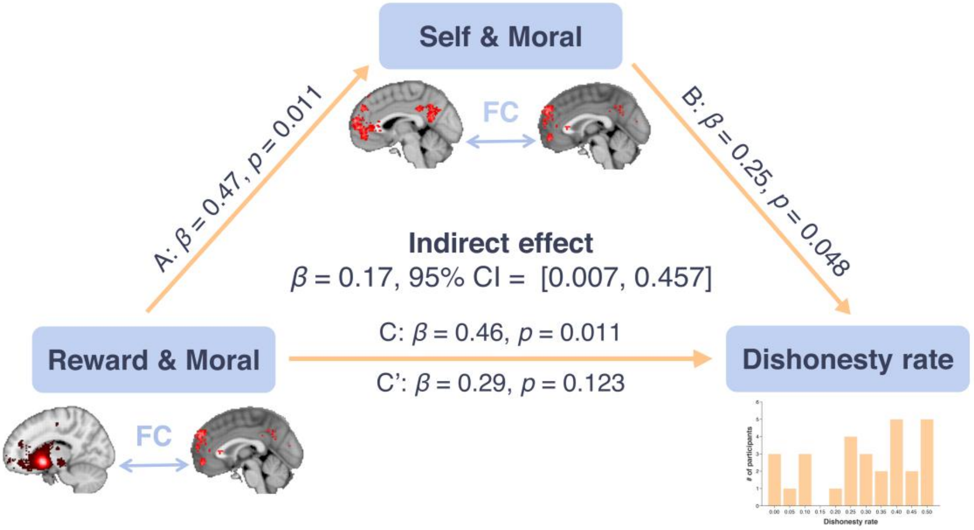


**Figure S7** *Mediation analysis of the self-referential network.* The functional connectivity between self-referential and moral networks could completely mediate the predictive effect of the functional connectivity between moral and reward networks on dishonesty rate.
